# Polymer dynamics of Alp7A reveals two ‘critical’ concentrations that govern dynamically unstable actin-like proteins

**DOI:** 10.1101/098954

**Authors:** Natalie A. Petek, Alan I. Derman, Jason A. Royal, Joe Pogliano, R. Dyche Mullins

**Author notes:** Corresponding author: Department of Cellular and Molecular Pharmacology, University of California, San Francisco, Genentech Hall, Room N312, Box 2200, 600 16th Street, San Francisco, CA 94158.

## Abstract

Dynamically unstable polymers capture and move cellular cargos in both bacteria and eukaryotes, but the regulation of their assembly remains poorly understood. Here we describe polymerization of Alp7A, a bacterial Actin-Like Protein (ALP) that segregates the low copy-number plasmid pLS20 in *Bacillus subtilis.* Purified Alp7A forms dynamically unstable polymers with two critical points: an intrinsic critical concentration (0.6 μM), observed when ATP hydrolysis is blocked, and a dynamic critical concentration (10.3 μM), observed when ATP hydrolysis occurs. From biochemical and kinetic analysis, the intrinsic critical concentration reflects a balance between filament elongation and shortening, while the dynamic critical concentration reflects a balance between filament nucleation and catastrophic disassembly. Although Alp7A does not form stable polymers at physiological concentrations, rapid nucleation by an accessory factor, Alp7R, decreases the dynamic critical concentration into the physiological range. Intrinsic and dynamic critical concentrations are fundamental parameters that can be used to describe the behavior of all dynamically unstable polymers.

## INTRODUCTION

Bacterial Actin-Like Proteins (ALPs) form polymers that participate in many biological processes, including: cell wall synthesis^1−4^, organelle positioning^5^, cytokinesis^6^, and DNA segregation. Although ALP molecules exhibit limited sequence homology with conventional actins, some ALP polymers share fundamental structural and biochemical features with their eukaryotic relatives, including a two-stranded filament architecture and an assembly mechanism that begins with formation of a trimeric nucleus. Because they are regulated by few accessory factors, bacterial ALPs can display more complex and varied assembly dynamics than eukaryotic actins, including: rapid spontaneous nucleation, a tendency to form filament bundles, and a dramatic decrease in polymer stability linked to hydrolysis of ATP.

The most well understood ALPs are encoded by Type-II DNA-segregation operons found on low copy-number plasmids. The simplicity of their cellular function makes DNA-segregating ALPs attractive model systems for studying how polymer assembly moves cellular cargo through the cytoplasm. Although different Type-II operons construct similar DNA-segregating structures, the molecular mechanisms underlying their assembly and function can differ significantly. For example, the two most studied DNA-segregating ALPs, ParM and AlfA, exhibit different polymerization dynamics, which are harnessed in different ways to capture and move DNA molecules. ParM filaments switch between stable growth and catastrophic disassembly ^7^, while AlfA polymers undergo a form of polarized growth and shortening known as treadmilling ^8^. Both catastrophic depolymerization ^9^ and rapid treadmilling ^10^ are also features of eukaryotic microtubules, linked to their inherent dynamic instability. Understanding how different ALP polymers switch between stable and unstable states, provides deeper insight into dynamic instability, and the ways in which it is harnessed to perform cellular functions.

In addition to an ALP, Type-II plasmid segregation also requires a centromeric DNA sequence and an adaptor protein that form a kinetochore-like complex called a segrosome. In the most thoroughly described systems this segrosome harnesses the free energy of polymer assembly to push the DNA cargo through the cytoplasm. Additionally, when two plasmids come into proximity, anti-parallel association of attached ALP filaments promotes formation of a bipolar ‘spindle’ between the plasmids, and elongation of ALP filaments within this spindle pushes the plasmid DNA molecules apart ^8,11,12^. In the present study we describe the self-assembly of a plasmid-segregating ALP, Alp7A, which shares less than 16% sequence identity with previously studied ALPs and whose in vivo behavior ^13^ poses challenges to the model of plasmid segregation derived from ParM and AlfA. The *alp7A* gene forms part of the *alp7AR* operon found in the pLS20 plasmid of *Bacillus subtilis* (natto). In addition to the actin-like protein Alp7A, this operon encodes Alp7R, a member of the Ribbon-Helix-Helix (RHH) family of dimeric DNA-binding proteins ^14^ The Alp7R dimers bind to nine nearby DNA sequence repeats (the *alp7C* locus), to form a DNA-protein complex that is required for plasmid partitioning ^13,15^. A previous study suggested that in vivo Alp7A forms dynamically unstable filaments that interact with the Alp7R/C complex to create a DNA-segregating spindle. In the absence of the Alp7R/C complex, however, polymer formation requires significant overexpression of Alp7A, and polymers formed at these high expression levels do not appear to be dynamic or filamentous^13^.Together, these observations suggest that Alp7A does not form filaments at normal cellular concentrations in the absence of the segrosome. This result contrasts sharply with the regulation of eukaryotic actins, whose cellular concentration greatly exceeds the critical concentration for filament assembly. Eukaryotes, therefore, require accessory factors such as profilin, thymosin-b4, and capping protein to maintain a large pool of monomeric actin far from its intrinsic steady state concentration.

In vitro we observe steady-state polymer formation only when the Alp7A concentration exceeds 10.3 μM, which is likely higher than the normal cellular concentration of this protein^1^. Surprisingly, ATP hydrolysis assays and experiments with hydrolysis-dead mutants reveal that Alp7A can form polymers at concentrations as low as 0.6 μM. The explanation for this apparent paradox is that filaments formed by Alp7A, and other dynamically unstable polymers, must satisfy two independent criteria for stability, each defined by a different critical concentration. The first stable point, which we call the intrinsic critical concentration, reflects a balance between filament elongation and shortening. This balance point is identical to the critical concentration that governs assembly of eukaryotic actin filaments ^16^. The second stable point, which we call the dynamic critical concentration, reflects a balance between the rates of filament nucleation and catastrophic loss. Because eukaryotic actin filaments disappear slowly, by diffusive rather than catastrophic processes, they do not exhibit this type of critical concentration. Thus, dynamically unstable ALPs do not require accessory factors to maintain a high monomer concentration, and regulatory factors that change the rate of filament nucleation can change the steady state polymer concentration by modulating the dynamic critical concentration —without necessarily affecting the intrinsic critical concentration. Consistent with this idea, we find that the dynamic critical concentration of purified Alp7A is sensitive to the rate of filament nucleation and can be modulated by lateral association of two-stranded filaments into stable bundles. The accessory factor, Alp7R, simultaneously accelerates Alp7A nucleation; inhibits bundling; and partially stabilizes isolated filaments. The net result of these different effects on polymer stability is to change the nature of the dynamic critical concentration from being the point at which bundles form to being the point at which nucleation is fast enough to outpace the rate of catastrophic filament loss. Our results reveals how accessory factors control the dynamic critical concentration of unstable actin-like filaments by multiple mechanisms that are independent of the underlying kinetics of polymer growth and shortening. Finally, our in vitro and in vivo experiments also reveal that Alp7R does not require DNA to nucleate and stabilize Alp7A filaments. Soluble Alp7R dimers are sufficient to increase Alp7A nucleation in vitro, and DNA-independent clustering of Alp7R is sufficient to drive filament formation in vivo. Based on similarities in operon architecture, we propose that this is a fundamental feature shared by other Type-II plasmid segregation systems.

## RESULTS

### The interaction of Alp7A with Alp7R: effects on nucleation and polymer architecture

Derman et al. (2009) found that co-expression of Alp7R was required to observe dynamic Alp7A polymer formation in *B. subtilis* cells. To understand the nature of this regulation, we purified Alp7A and Alp7R and studied their interaction in vitro. Addition of 5 mM Mg-ATP to 15 μM apo-Alp7A induced a time-dependent increase in right-angle light scattering, consistent with a nucleation-limited polymerization reaction. The initial rate of change in light scattering intensity was slow, but subsequently accelerated before slowing to a final steady-state plateau by approximately 500 seconds (Figure 1A, green trace). Kinetic analysis of these light scattering data, based on the work of Nishida and Sakai ^17^, suggests that formation of a stable Alp7A nucleus requires four kinetically resolvable steps (Figure 1B, green trace). Addition of 0.5 μM Alp7R dramatically increased the rate at which 15 μM Alp7A polymerized, and reduced the time required to reach steady state to approximately 40 seconds (Figures 1A, red trace). Alp7R also decreased the number of steps required to form a nucleus from four to one (Figure 1B, red symbols), suggesting that it accelerates polymerization at least partly by accelerating nucleation. Based on the steady-state light scattering signal we measured a critical concentration of 10.3 μM for the assembly of Alp7A polymers, which evidence suggests ^1^ is above the cellular concentration of Alp7A. Addition of increasing amounts of Alp7R decreases the apparent critical concentration of Alp7A polymerization (Figure 1C-F), down to a minimum value of approximately 2 μM in the presence of 5 μM Alp7R (Figure 1F). Above 5 μM, however, Alp7R causes the apparent critical concentration of Alp7A to rise, back toward its original value in the absence of Alp7R. This biphasic effect on polymerization suggests that the activity of an Alp7R dimer relies on its interaction with at least two Alp7A monomers. At high concentrations, Alp7R dimers are less effective because they compete for a limited pool of Alp7A monomers. Surprisingly, the effects of Alp7R on nucleation and critical concentration of Alp7A filaments do not require the presence of DNA.

**Figure 1:**
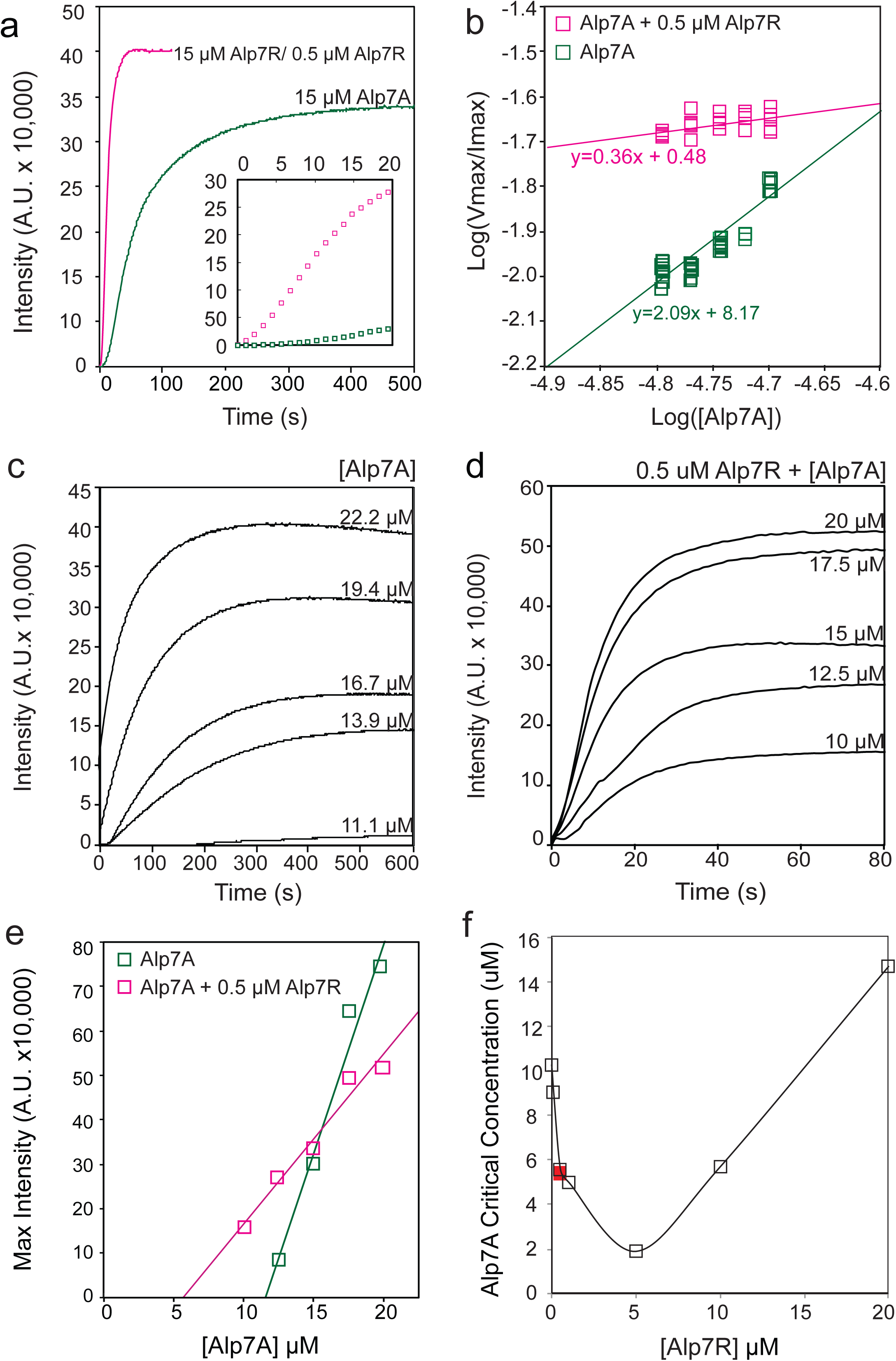
Alp7R dimers enhance nucleation and decrease critical concentration for Alp7A polymer formation. (a) Right-angle light scattering signal of 15 μM ATP-Alp7A polymerizing in the presence (pink) or absence (green) of 0.5μM Alp7R. Inset shows a zoom of the first 20 seconds of the reaction following the addition of 3mM ATP/Mg++. Buffer conditions: 25 mM Tris pH 7.5, 100 mM KCl, 1 mM Mg++Cl2, 1 mM DTT. (b) Analysis of the nucleation mechanism by the method of Sakai et al. wherein the log of the maximum velocity is plotted versus the log of protein concentration. Twice the slope of the line plotted through these points describes the number of kinetically resolvable nucleation steps. (c) Polymerization of increasing concentrations of Alp7A in the presence of 3 mM ATP/Mg++ visualized by right angle light scattering. (d) Titration of Alp7A in the presence of 0.5 μM Alp7R. (e) Maximum intensity plots of the titrations of Alp7A from ‘c’ and ‘d,’ green squares and pink squares respectively. The x-intercept of these lines represents the critical concentration of Alp7A for each condition. (f) A plot of the critical concentration of Alp7A as a function of Alp7R concentration. The red square represents the critical concentration of Alp7A in the presence of 0.5 uM Alp7R/50 nM alp7C.

In addition to accelerating filament assembly, Alp7R also altered the architecture of Alp7A polymers. By electron microscopy, Alp7A appears as ribbon-like bundles of two-stranded filaments (Figure 2A), but addition of Alp7R reduced the size of Alp7A bundles in a concentration-dependent manner (Figure 2B). Low concentrations of Alp7R (1:20 stoichiometry with Alp7A) had little effect on bundle size, while higher Alp7R concentrations (1:2 stoichiometry with Alp7A) converted the Alp7A bundles into well separated, two-stranded polymers (Figure 2A-B, S1A), similar in size to eukaryotic actin filaments. As with its effect on filament nucleation, the ability of Alp7R to antagonize Alp7A bundle formation did not depend on the presence of DNA (Figures 1F, 2C).

**Figure 2:**
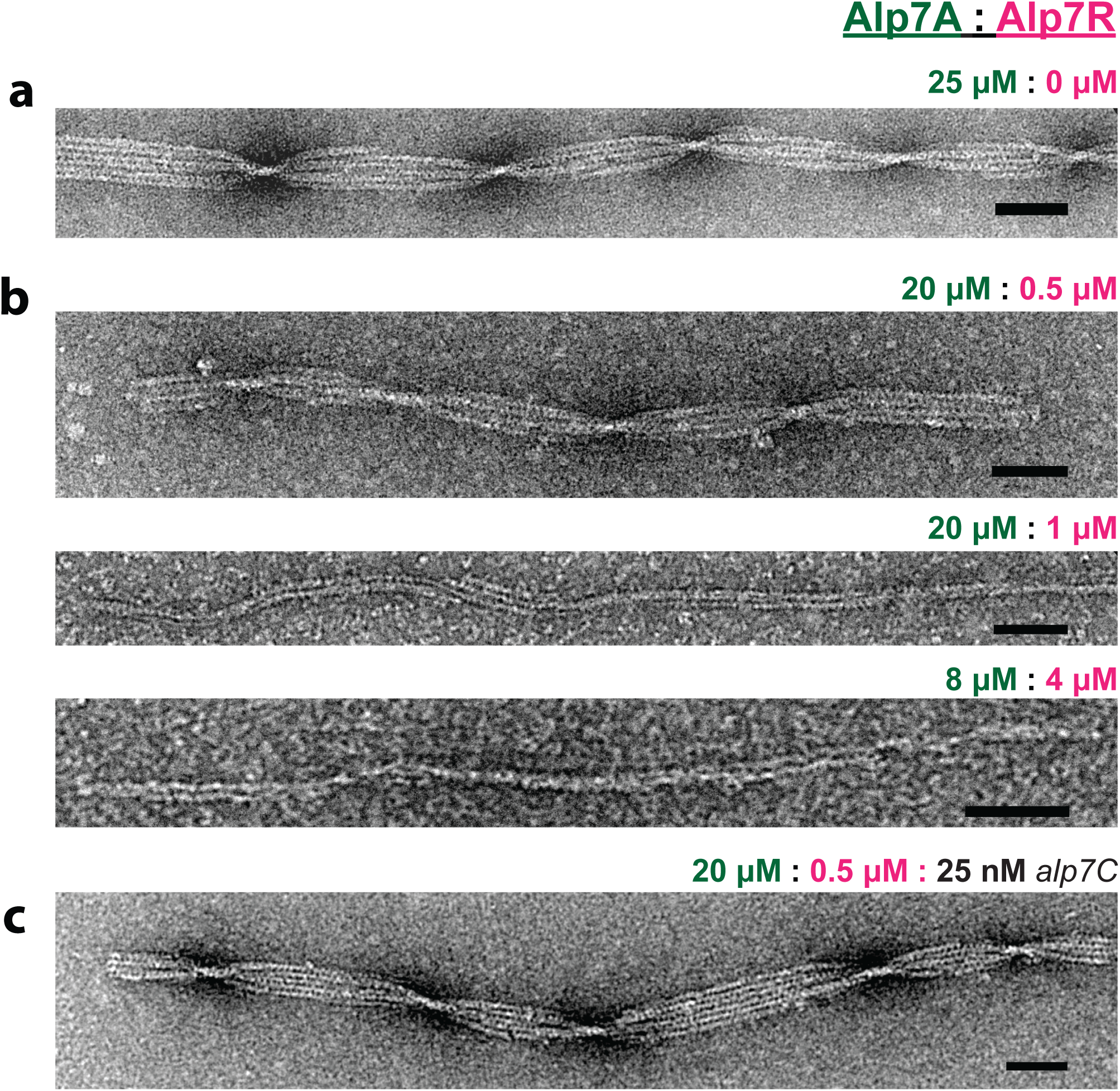
Alp7R disrupts bundling of Alp7A. Electron micrographs of Alp7A polymers adsorbed to carbon-coated grids and stained with 0.75% uranyl formate. (a) Alp7A alone at a final concentration of 20 μM. (b) Alp7A polymerized in the presence of Alp7R: (top) 20 μM Alp7A with 0.5 μM Alp7R, (middle) 20 μM Alp7A with 1.0 μM Alp7R, and (bottom) 8 μM Alp7A with 4 μM Alp7R. (c) 20 μM Alp7A, 0.5 μM Alp7R 50 nM alp7C DNA. For all images, buffer conditions same as Figure 1, polymerized with 3 mM ATP/Mg++. In all panels Alp7A polymerization was induced by addition of 3 mM ATP/Mg++. Scale bars 25 nm.

### Alp7R does not require DNA to nucleate filaments in vivo

Since Alp7R did not require DNA to enhance Alp7A polymerization or block bundle formation in vitro, we tested whether DNA binding is required for Alp7R to promote Alp7A filament formation in vivo. In addition to specific binding sites within *alp7C*, Alp7R also binds weakly to non-specific DNA. When expressed by itself, Alp7R-GFP localizes to the chromosome, similar to DAPI-staining (Figure 3A). To prevent Alp7R from interacting non-specifically with DNA, we tethered the protein to the cell envelope by fusing the *alp7R* gene to the divIVA gene, which encodes a cell division protein that is assembled into ring-shaped structures at the cell poles and septa ^18^. To gauge the effectiveness of this strategy, we expressed a *divIVA-alp7R-gfp* gene fusion, which gave rise to the polar and septal rings characteristic of DivIVA (Figure (3B). We observed no evidence of interaction with chromosomal DNA, or interference with DivIVA assembly. We next expressed *divIVA-alp7R* together with *alp7A-gfp*. In this case we observed Alp7A-GFP localized to septal and polar rings and dynamic filaments emanating from these static rings (Figure 3D-E; M1).

**Figure 3:**
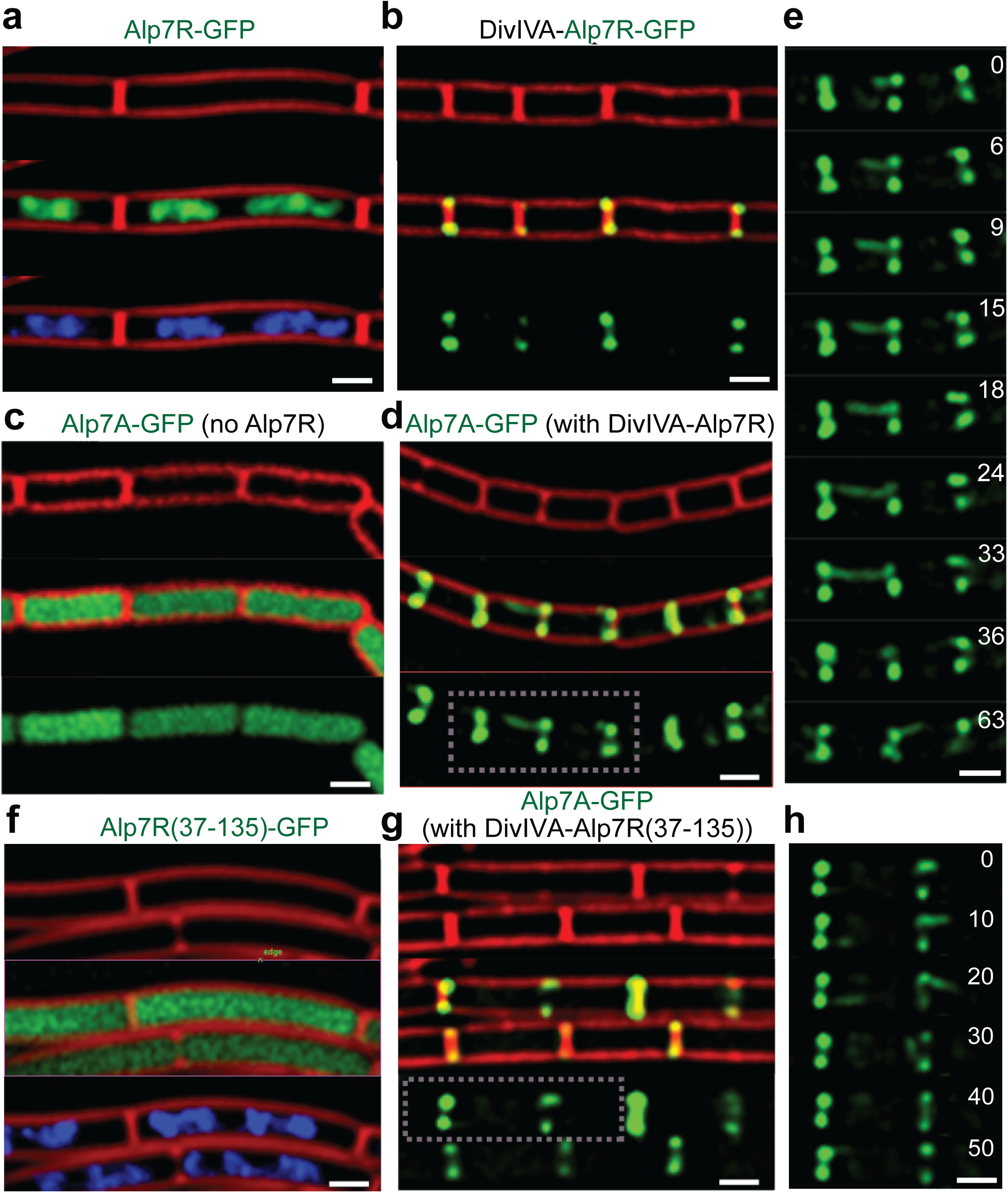
Alp7R nucleates filaments in vivo in the absence of DNA. Live-cell fluorescence microscopy of B. subtilis expressing fluorescent derivatives of Alp7A or Alp7R together with fluorescent markers for membranes and DNA. (a) Alp7R-GFP nonspecifically coats DNA in the absence of the specific alp7C DNA sequence. The top panel shows membrane in red, middle is Alp7R-GFP, bottom shows DNA by DAPI staining. (b) The DivIVA-Alp7R fusion localizes to the cell pole as seen by: membrane staining (top) overlay (middle) and DivIVA-Alp7R-GFP (bottom). (c) Alp7A-GFP is diffuse in the absence of Alp7R: cell membranes (top), overlay (center), Alp7A-GFP (bottom). (d) Alp7A forms filaments that nucleate from the cell pole in the presence of the DivIVA fusion protein. The top panel shows cell membranes, the center shows the overlay, and the bottom panel shows filaments of Alp7A-GFP nucleating from the cell pole. (e) A montage of two cells (dotted square in d) over time shows Alp7A-GFP nucleating, elongating and undergoing catastrophic disassembly (left cell). All scale bars: 1 μm.

To rule out a specific requirement for DNA binding, we constructed a variant fusion protein with a disrupted DNA-binding domain. The resemblance of Alp7R’s amino-terminal region to DNA-binding proteins of the RHH2 family together with the domain organization of similar proteins, such as ParR, suggested that the DNA-binding domain of Alp7R is at its amino terminus. We therefore removed the first 36 codons from *alp7R* in both the *divIVA-alp7R* and *divIVA-alp7R-gfp* contexts, to abolish DNA binding. Alp7R(37-135)-GFP failed to localize to the chromosome as previously seen for wild type Alp7R^2^, but rather distributed uniformly throughout the cytoplasm (Figure 3F). The DivIVA-Alp7R(37-135)-GFP fusion protein produced DivIVA-like polar and septal rings (Figure 3G), and expression of *divIVA-alp7R(37-135)* together with *alp7A-gfp* gave rise to the same profile of the original *divIVA-alp7R* fusion, with dynamic filaments emanating from septal and polar rings (Figure 3G, 3H). In this case though, filaments formed with no possibility of DNA binding, consistent with our biophysical finding that Alp7R could nucleate Alp7A filament formation independently of DNA in vitro.

### ATP hydrolysis and Alp7A polymer assembly

Consistent with polymerization-induced hydrolysis of ATP, the steady-state rate of phosphate release by ATP-Alp7A increased non-linearly with protein concentration (Figure 4A). At low concentrations of Alp7A phosphate release was undetectable, but beginning around 1 μM the rate of phosphate release increased rapidly. Barring ‘action at a distance’ this concentration-dependent increase in specific ATPase activity of Alp7A must reflect self-association of the protein into oligomeric species. In addition, individual phosphate release traces (Figure 4B) reveal multi-phase kinetics of Alp7A oligomerization below the apparent critical concentration. Addition of ATP to 10 μM Alp7A, for example, induces a rapid burst of ATP hydrolysis that slows to a steady state rate at about the same time that polymerization of low concentrations of Alp7A reaches a steady-state plateau in light-scattering assays (Figure 1C).

**Figure 4:**
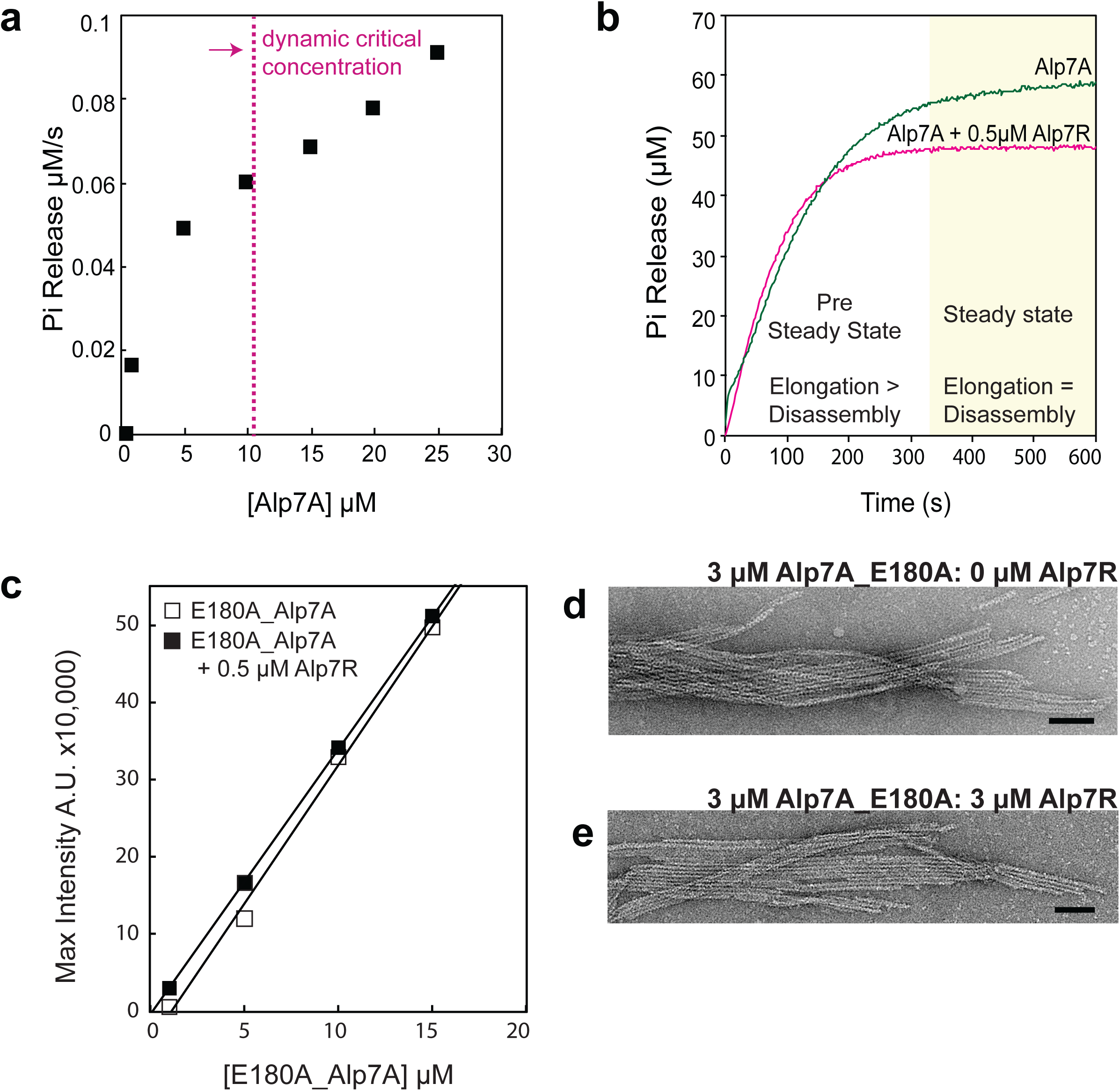
ATP hydrolysis reveals ephemeral filaments below the apparent Alp7A critical concentration which are stabilized by Alp7R. (a) The steady-state rate of phosphate release by Alp7A, measured 250-300 seconds after addition of ATP, increases non-linearly with Alp7A concentration. Below 0.5 μM Alp7A, phosphate release is undetectable, but rises rapidly above 0.5 μM, indicating the presence of polymer. Above 5 μM Alp7A, the slope decreases, suggesting that monomer dissociation has become rate limiting for the formation of new polymer. (b) The biphasic kinetics of individual phosphate release traces reveal the assembly dynamics of 10 μM Alp7A, below its apparent critical concentration. Similar initial rates of phosphate release are observed in the absence and presence of 0.5 μM Alp7R, but at steady state (after 300 seconds) Alp7R slows the rate of phosphate dissociation, suggesting that it slows the rate of filament turnover. Buffer conditions same as Figure 1, polymerized with 10 mM ATP/Mg++. (c) The hydrolysis-dead mutant Alp7A_E180A has a critical concentration of 0.6μM Alp7A (open squares), which is slightly reduced in the presence of Alp7R (closed squares). (d) 3μM Alp7A_E180A forms bundled sheets (top) that are not disrupted by addition of Alp7R (bottom). Buffer conditions same as Figure 1, polymerized with 3 mM ATPMg++.

From steady-state rates of phosphate release measured below 10 μM, we estimate that less than 0.15 μM Alp7A is present in oligomeric species below the apparent critical concentration. Assuming that ATP hydrolysis occurs only on trimers and larger species, we estimate the concentration of ATP-hydrolyzing oligomers to be less than 0.05 μM. Although the presence of ~10 μM monomeric Alp7A (a 220-fold excess) posed a challenge, we investigated these oligomeric species by electron microscopy and dynamic light scattering. Although consistent with the result with the ATPase measurements, neither of these techniques provided insight into the nature of these small oligomers due to the high signal to noise ratio.

Above 5 μM Alp7A, the steady-state rate of phosphate release decreased, suggesting that in this regime monomer dissociation becomes rate limiting for the formation of new polymer (Figure 4A). Addition of Alp7R further decreased the steady state rate of ATP hydrolysis by Alp7A in this regime (Figure 4B), suggesting that Alp7R promotes polymer formation at these concentrations, at least in part, by reducing the rate of disassembly of these ephemeral Alp7A filaments.

To better understand the role of ATP hydrolysis in the assembly and disassembly of Alp7A polymers, we constructed a hydrolysis-defective mutant of Alp7A. From our crystal structure, we identified a residue in the nucleotide binding pocket of Alp7A (E180) analogous to residues in ParM (E148) and actin (Q137) required for ATP hydrolysis^7,19^ We previously showed that this mutant forms stable filaments at low concentration in vivo and hypothesized that it was due to a hydrolysis defect. We found that Alp7A-E180A forms stable filaments with an apparent critical concentration much lower (0.6 ± 0.4 μM) than that of wild-type Alp7A (Figure 4C). This concentration matches the concentration at which we first observe evidence of ATP hydrolysis by wild type Alp7A, indicating that it reflects a ‘critical’ concentration for both the wild type and hydrolysis defective Alp7A mutant. Addition of Alp7R, however, does not significantly decrease the critical concentration for assembly of the E180A mutant (Figure 4C). We propose, therefore, that this value represents an intrinsic critical concentration for Alp7A filaments, analogous to the ATP critical concentration of eukaryotic actin, which is set by the rates of filament elongation and shortening.

The hydrolysis-defective mutant also forms bundles, which are not disrupted by addition of Alp7R (Figure 4D), suggesting that the ability of Alp7R to disrupt bundle formation requires ATP hydrolysis and filament turnover. By contrast to the wild-type Alp7A, the bundles of the E180A mutant is stable in high salt. Attempts to disrupt the lateral filament associations with 1.5M NaCl instead resulted in a large fraction of the polymer appearing as three-dimensional bundles rather than flattened sheets (Figure 5A-B). This is likely because high salt weakens the interaction between Alp7A bundles and the negatively charged surface of the carbon-coated grid^20^, better preserving their three dimensional structure. Consistent with the idea that flattened ribbons are an artifact of interaction with the carbon-coated EM grid, when we diluted filaments in water we observed only flattened sheets. We are unable to see this result using the wild-type Alp7A, because dilution of these filaments results in their rapid and complete disassembly (Figure 5C).

**Figure 5:**
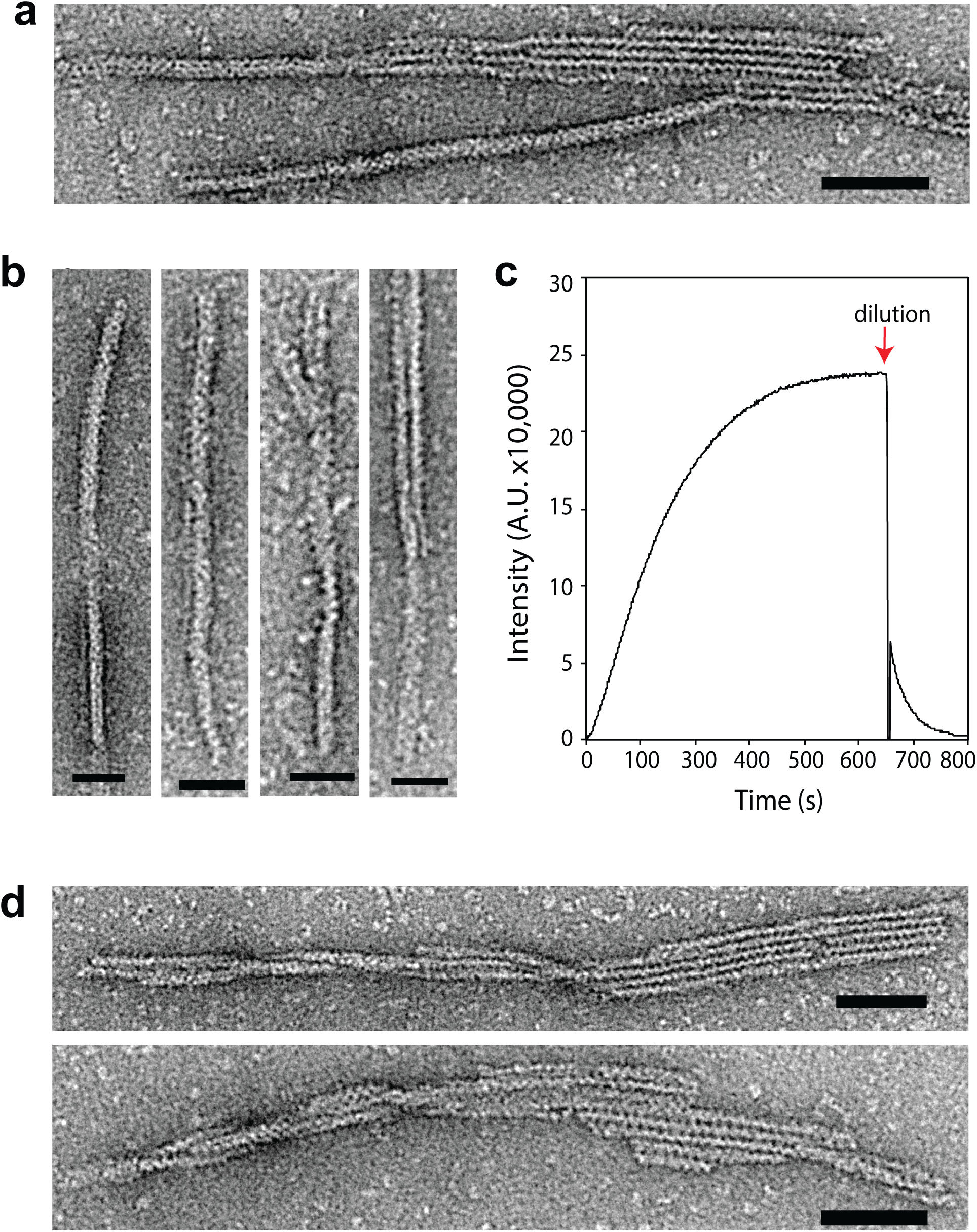
Flat Alp7A ribbons form more compact bundles in high salt. Electron micrographs of Alp7A polymers formed in different concentrations of NaCl; adsorbed to carbon-coated grids; and stained with 0.75% uranyl formate. (A-b) Addition of 1.5M NaCl does not disassociate Alp7A bundles, but instead partly shields the bundles from the surface charge of the carbon-coated grid, diminishing the occurrence of flattened sheets. High salt bundles are 10-15nm in width, which are likely one to two-stranded bundle. (c) Rapid disassembly of polymerized Alp7A at steady state. At the red arrow 25uM Alp7A is diluted 3x by addition of polymerization buffer; final Alp7A concentration is 8.3 μM, and ATP is 3.3 mM. (d) E180A_Alp7A filaments diluted in 0.5M NaCl. Filaments are both flattened sheets and twisted filament bundles. Scale bars: B = 25nm. A and D = 50nm. Buffer conditions same as Figure 1, polymerized with (a-b, d) 3mM ATP/Mg++ and (c) 10 mM ATP/Mg++.

**Figure 6:**
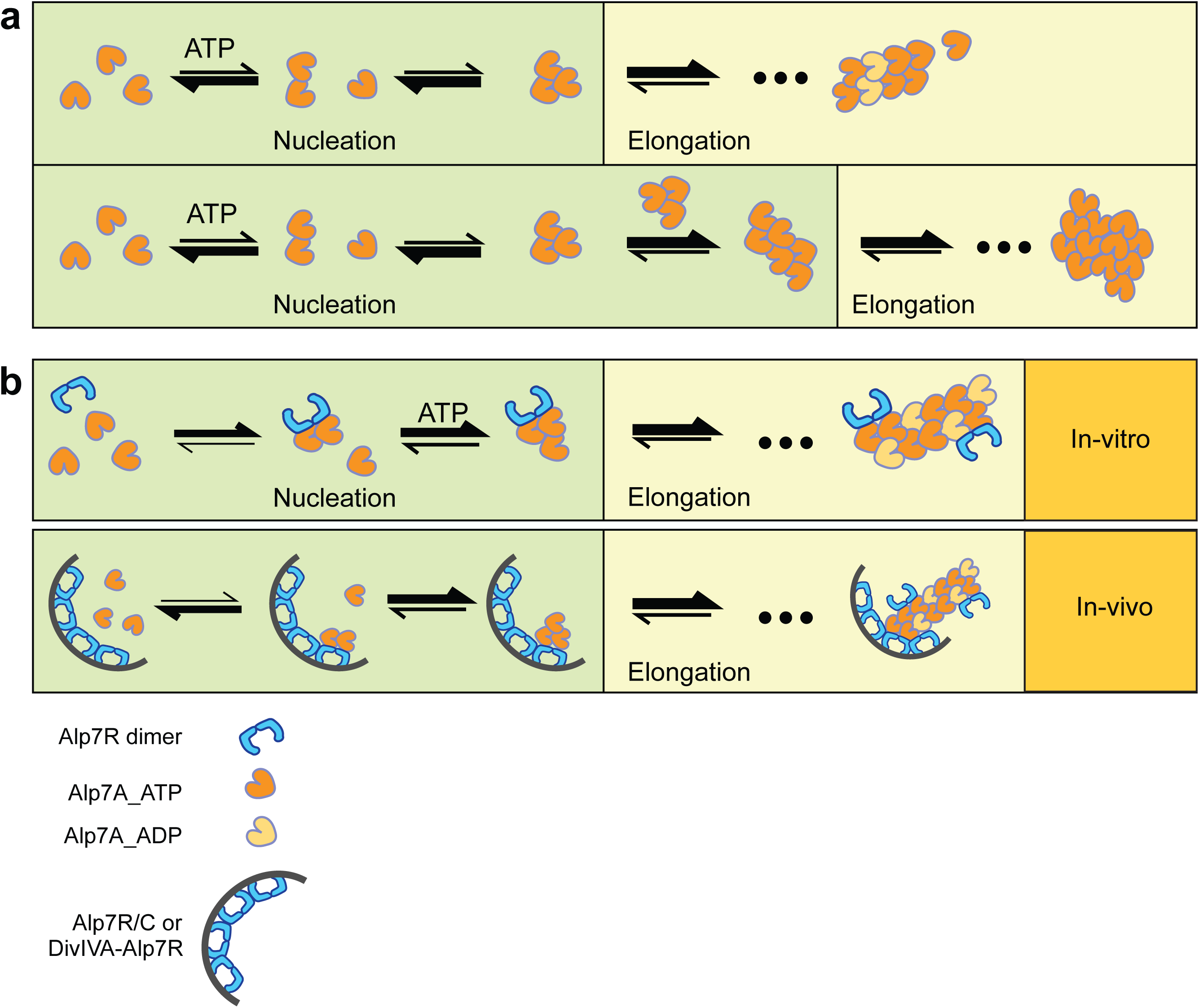
Assembly of Alp7A polymers and regulation by Alp7R. (a) Nucleation of a simple, two-stranded actin-like filament (top) requires fewer steps than nucleation of Alp7a polymers, which require lateral filament association for stability (bottom). The sizes of the bi-directional arrows reflect the relative rates of association and dissociation of monomers from various oligomers and polymers. (b) Alp7R dimers alter the assembly mechanism of Alp7A by promoting rapid nucleation and antagonizing bundle formation. In our in vitro experiments (top) Alp7R dimers bind two Alp7A monomers in a “pre-nucleation” complexes. Upon addition of ATP, only one kinetically resolvable step is required to form a stable nucleus. Side-binding of Alp7R to Alp7A filaments antagonizes bundle formation and slows the rate of filament disassembly. In vivo (bottom), Alp7R dimers are clustered by their interaction with tandem DNA sequence repeats in the alp7C locus, providing a higher avidity interaction with Alp7A filaments.

## DISCUSSION

Our current understanding of plasmid-segregating ALPs began with structure-function studies of the R1 par operon, and has since expanded to include comparisons of the molecular mechanisms employed by an evolutionarily diverse set of systems ^21−23^. The three best-studied ALPs —ParM, AlfA, and now Alp7A— share two properties that appear to define minimal requirements for plasmid segregation: (i) polymer destabilization driven by intrinsic ATPase activity and (ii) polymer stabilization caused by anti-parallel bundling and plasmid association. The very low sequence conservation among these ALPs and the significant divergence of other structural and biochemical features suggest that these shared properties are essential for plasmid segregation and have been maintained by positive selection.

### Multiple mechanisms for stabilizing dynamic filaments

Apart from a few shared properties, plasmid-segregating ALPs exhibit remarkable diversity in assembly dynamics and regulation. For example, the polymerization of different ALPs is initiated in different ways. ParM has an extremely high rate of spontaneous nucleation and ParR/parC segrosome may simply capture and stabilize preexisting filaments. AlfA has a much slower rate of spontaneous assembly. Rapid nucleation of AlfA polymers requires a segrosome that contains the accessory protein, AlfB, and the centromeric sequence, parN, and that is sensitive to the stoichiometry of this AlfB:parN complex. In contrast to both ParM and AlfA, spontaneous assembly of Alp7A filaments is relatively slow and requires only the accessory protein Alp7R for rapid nucleation. Plasmid movement must, therefore, rely on the clustering of multiple Alp7R molecules on the *alp7C* locus to maintain contact between the DNA and the growing polymer. Such an effect would be similar to the way clustering of eukaryotic actin polymerases, such as VASP, promote processive interaction with the growing ends of actin filaments ^24−26^.

The key to understanding the common principles that transcend this mechanistic diversity lies in the relationships between four fundamental parameters, the rate constants that describe filament: (i) elongation, (ii) shortening, (iii) nucleation, and (iv) catastrophe. Together, the rates at which ATP-bound filaments elongate and shorten define an intrinsic critical concentration, above which individual filaments grow steadily. Wild-type eukaryotic actin and mutant ALPs that are defective in ATP hydrolysis do not experience catastrophic depolymerization and so their assembly is dominated by this intrinsic ATP critical concentration^3^. For wild-type ALPs the balance between rates of filament creation (nucleation) and loss (catastrophe) defines a second, or dynamic, critical concentration, above which the steady-state number of filaments is greater than zero. The dynamic critical concentration of ParM, AlfA, and Alp7A is all higher than their intrinsic critical concentration but this value depends on the propensity of the filaments to form bundles and it can be shifted up or down by accessory factors. Interestingly, ParM has a high rate of spontaneous nucleation and a weak tendency form bundles, so its dynamic critical concentration is determined primarily by the rate of filament loss due to dynamic instability. In contrast, Alp7A has a much lower rate of spontaneous nucleation, which elevates the concentration of Alp7A at which nucleation balances catastrophe to a value much higher than that of ParM. Alp7A, however, has a strong, tendency to form stable bundles, so its dynamic critical concentration is set by the rate at which newly formed filaments interact with each other. In this case, Alp7A polymers appear when the average filament lifetime becomes comparable to the time required for multiple filaments to associate into a stable bundle. The role of bundling in this monomer-to-polymer transition can be seen in the large number of steps required to form stable Alp7A bundles. Addition of the accessory factor, Alp7R, simultaneously increases the rate of Alp7A nucleation, stabilizes filaments against disassembly, and disrupts bundling, making Alp7A filaments behave much more like ParM filaments. This result suggests that, while anti-parallel bundling is important for making a bipolar spindle ^8,12^, the stability of these bundles must be maintained within a window that is neither too high nor too low for proper spindle assembly and function.

### DNA-independent effects of Alp7R on Alp7A polymerization

One surprising result of our work is that the accessory factor, Alp7R, does not require DNA for its effects on Alp7A filament assembly. This is surprising because the DNA binding component of a Type II plasmid segregation system must somehow couple actin-like filament assembly to the movement of DNA. Our in vitro and in vivo results both argue the *alp7C* locus performs this coupling function by simply clustering multiple Alp7R molecules on a small DNA locus. While soluble Alp7R induces rapid Alp7A nucleation throughout the cytoplasm, the Alp7R cluster at the segrosome provides a high-valency site of filament binding and stabilization. Therefore, despite significant differences in underlying assembly dynamics, both Alp7A and ParM filaments create DNA-segregating spindles with the same combination of fast, spatially distributed nucleation; localized filament stabilization; and weak anti-parallel bundling. Our results reveal that, while DNA segregation constrains parameters governing assembly of ALP-based spindles, the parameters themselves (e.g. rate of nucleation and strength of anti-parallel bundling) can either be intrinsic to the ALP or can be set by accessory factors.

Kinetic analysis reveals that Alp7R promotes single-step initiation of Alp7A polymers. In the simplest model for nucleation, Alp7R, which binds Alp7A monomers and forms dimers {{Derman:2012ko}, and this study}, brings two Alp7A protomers into contact in an orientation the mimics the end of a growing filament. The most parsimonious explanation for why Alp7R antagonizes formation of Alp7A bundles is that the Alp7R binding site responsible for nucleation remains accessible after monomers have assembled into a filament. To test the plausibility of this explanation we compared the crystal structures of Alp7A (from the companion study) to the structure of ParM bound to its regulatory peptide from ParR. Assuming that the Alp7A/Alp7R interaction is structurally similar to that of ParM/ParR, our data suggest that the interaction interface is accessible within the Alp7A filament.

## MATERIALS AND METHODS

### Expression and Purification of Alp7A

Alp7A was codon optimized (Mr. Gene) and cloned into the pet28a vector using the NcoI and HindIII restriction sites. Alp7A in Pet28 was freshly transformed for each purification into C43 cells and grown in TPM media at 37°C to an OD of 0.4-0.6, then shifted to 18°. Cultures were induced overnight with 0.2mM IPTG, then pelleted and washed with 1x PBS. Washed pellets were flash frozen in liquid nitrogen and stored at −80°C.

To purify native Alp7A, three liters of expression culture were resuspended in Lysis buffer (25mM Tris pH 7.6, 100 mM KCl, 1mM EDTA, 1mM DTT, PMSF). Cells were dounce homogenized prior to lysis by 3x passes through an Emulsiflex-C3 (Avestin). The lysate was cleared with a high-speed spin (60 min at 170,000×g, 4°C) to remove cellular debris and insoluble protein. Ammonium sulfate was added slowly to the cleared lysate to 50% solubility, lysate was slowly stirred at 4°C for one hour after all ammonium sulfate has been added. The precipitate is removed by centrifugation at 170,000xg for 30 minutes, 4°C. Alp7A remained soluble; the supernatant was therefore transferred to a clean ultracentrifuge tube and brought to room temperature using an ambient water bath. 5 mM ATP/6 mM MgCl_2_ were then added to induce polymerization. After a 15 minute incubation, the polymer was pelleted by high speed centrifugation (20 min at 170,000×g, 25°C) and then resuspended in 1/10 volume of cold depolymerization buffer (25mM Tris pH 7.6, 200mM KCl, 5mM EDTA, 1mM DTT). The resuspended protein was dialyzed overnight to remove residual ATP and ensure complete depolymerization. Alp7A was gel filtered on Superdex S-200 resin into polymerization buffer (25 mM Tris pH 7.6, 100 mM KCl, 1 mM MgCl_2_, 1mM DTT); the purest protein fractions were pooled and dialyzed into storage buffer (polymerization buffer + 20% glycerol). Aliquots were snap frozen in liquid nitrogen and stored at −80°C. Protein is quantified at A280, by the Alp7A extinction coefficient 34,965 M^-1^cm^-1^.

### Expression and Purification of Alp7R

A pE-SUMOstar vector (LifeSensors) containing the Alp7R insert was transformed into BL21 cells. Cultures were grown in LB media to an OD of 0.4-0.6, then induced for 4 hours with 0.750 mM IPTG. Cultures were pelleted and washed with 1x PBS. Washed pellets were flash frozen in liquid nitrogen then stored at −80°C.

To purify 6x-his-SUMO-Alp7R, cells from two liters of culture were resuspended in lysis buffer (50mM HEPES pH 8.0, 400mM NaCl, 1mM BME + cOmplete EDTA free protease inhibitor-Roche). Cells were dounce homogenized prior to lysis by 3x passages through an Emulsiflex-C3 (Avestin). The lysate was cleared with a high-speed spin (60 min at 170,000×g, 4°C) to remove cellular debris and insoluble protein. Protein was bound to a 5mL HiTrap Chelating HP column (GE Healthcare) precharged with CoCl_2_, washed with lysis buffer, then eluted with 500mM Imidazole. The most concentrated fractions were pooled, then incubated for 30 min at room temperature with SUMO Protease I (90+% cleaved after this step). This was then dialyzed over night in 50mM HEPES pH 8.0, 400mM NaCl, 1mM BME. The cleaved tag and uncleaved protein were then removed by running the protein over a 1ml HiTrap Chelating HP column. The eluate was gel filtered on Superdex S-200 resin. Peak fractions were pooled and dialyzed into 25mM HEPES, 400mM NaCl, 1mM DTT, 20% glycerol. Aliquots were snap frozen in liquid nitrogen and stored at −80°C. Protein was quantified by Sypro Ruby (S-12000. Molecular Probes^™^) staining as Alp7R contains no endogenous tryptophans.

### Amplification of *alp7C*

*alp7C* was amplified by PCR of the *alp7C* region from the mini-plS20 plasmid (courtesy of Joe Pogliano) using the methods described in Derman et. al., 2012.

### Negative Staining and Electron Microscopy

Samples were prepared by applying 4ul of polymerization mix to 200 mesh Formvar carbon film coated (Electron Microscopy Sciences-FCF200-Cu) glow discharged grids and incubating for 30 seconds. Grids were washed 3x in buffer, blotting between washes and then stained 3x with 0.75% uranyl formate, again blotting between applications. Micrographs were taken with a Technai T12 microscope using an acceleration voltage of 120 kV and a magnification of x27,000 or x48,000. Images were recorded with a Gatan 4k x 4k charge-coupled device camera.

### Right-Angle Light Scattering

Light scattering experiments were performed using an ISS K2 fluorimeter at a wavelength of 320nm. Polymerization of Alp7A was induced by hand mixing or use of a SFA-20 rapid mixer (Hi-Tech) coupled to the fluorimeter. For each condition four to five traces were background subtracted and averaged. The critical concentration, or steady state monomer concentration, of Alp7A was determined by plotting the maximum light scattering intensity of averaged traces versus the concentration of Alp7A. The x-intercept of this line is the value for the critical concentration.

### Plasmids and plasmid construction

The plasmids used in this study are listed in Supplementary Table 1. The construction of plasmid pAID3219 (pP_xyl_alp7R-gfp), where gfp encodes green fluorescent protein, was described previously (Derman et al., 2012). Plasmid pAID3604 (pP_xyl_alp7RΔ36-gfp) was constructed by amplification of pAID3129 with oligonucleotide primers P1 and P2, restriction of the amplicon with AvrII and SphI, and ligation to pAID3129 restricted with AvrII and SphI. Plasmids and ligation mixtures were introduced into E. coli DH5α by electroporation or by transformation of chemically competent cells.

### Bacterial strains, strain construction, and growth of bacteria

The strains used in this study are listed in Supplementary Table 1. Strain JP3348 (PY79 amyE::P_hyspank_divIVA-alp7R-gfp) was constructed by integration into the PY79 chromosome of a modified version of plasmid pDR197, a derivative of the *B. subtilis* integration vector pDR111[31] that contains a divIVA-gfp gene fusion under control of the hyperspank promoter (Ramamurthi and Losick. 2009). Plasmid pAID3129 was amplified with oligonucleotide primers P3 and P4, and the amplicon was cloned into the pCR2.1-TOPO vector (Life Technologies). The 429 bp segment containing the alp7R gene followed by sequence coding for the seven amino acid linker RPEDIH was excised with NheI and ligated to pDR197 restricted with NheI. The tripartate gene fusion on the resulting plasmid was then integrated into the PY79 chromosome at amyE by a double recombination event. The same strategy was used to construct strain JP3350 [PY79 amyE::P_hyspank_divIVA-alp7R_TAA_-gfp thrC::(P_xyl_alp7A-gfp)], except that primer P5, which preserves the alp7R stop codon, was used in place of P2, and strain JP3161 [PY79 thrC::P_xyl_alp7A-gfp)] was used for integration. Strains JP3569 [PY79 amyE::(lacI^+^ P_hyspank_divIVA-alp7RΔ36-gfp spec)] and JP3568 [JP3161 amyE::(lacI^+^ P_hyspank_divIVA-alp7RΔ36_TAA_-gfp spec)] were constructed by integration into PY79 and JP3161 of versions of the integration plasmids in which the first 36 codons of alp7R had been deleted. The alp7RΔ36 deletion in JP3569 was constructed by amplification of the P_hyspank_divIVA-alp7RΔ36-gfp integration plasmid with oligonucleotide primers P6 and P7, restriction of the amplicon with NheI and AatII, and ligation to pDR197 restricted with NheI and AatII. The alp7RΔ36_TAA_ deletion in JP3568 was constructed by oligo-directed deletion mutagenesis of the P_hyspank_divIVA-alp7RΔ36_TAA_-gfp integration plasmid with mutagenic oligonucleotide primers P8 and P9 as described in Derman et al., 2012. These plasmids were then integrated into PY79 or JP3161 at amyE. The orientation of each alp7R insertion was evaluated by restriction endonuclease digestion and the sequences verified by DNA sequencing. Only background green fluorescence could be detected from control strains JP3349 [PY79 amyE::P_hyspank_divIVA-alp7R_TAA_-gfp] and JP3567 [PY79 amyE::P_hyspank_divIVA-alp7RΔ36_TAA_-gfp] at all induction levels surveyed, indicating that there was no read-though into gfp when the alp7R stop codon was present. Media and antibiotic supplementation was as described in Derman et al., 2012.

### Live-Cell Microscopy

Agarose pads for microscopy were prepared and imaged as described in Derman et al., 2012.

### Oligonucleotide Primers

P1: 5’-gtactcctagggaggcacaatcaatttaggtgattagaaatgaaaagtaagggtacctttagagagtatgc-3’

P2: 5’-gtactgcatgcgacctcgtttccaccggaattag-3’

P3: 5’-ctgtagctagcatggggaaaaacaaaagaattccac-3’

P4: 5’-gtactgctagcatgtatatctccttccggccgaaaatcatagtcgtattcttcttcaatag-3’

P5: 5’-gtactgctagcatgtatatctccttccggccgttaaaaatcatagtcgtattcttcttcaatag-3’

P6: 5’-ctgtagctagcagtaagggtacctttagagag-3’

P7: 5’-catttatcagggttattgtctc-3’

P8: 5’-gaggaaaaggaagctagcagtaagggtacctttagagag-3’

P9: 5’-ctctctaaaggtacccttactgctagcttccttttcctc-3’

### Phosphate Release Assays

Phosphate release assays were performed using the Enzcheck Phosphate Assay Kit (Invitrogen-E6646) and an Ultraspec 2100 Pro spectrophotometer (GE Healthcare Lifesciences). Both protein, and ATP/MgCl2 mixtures were pre-incubated with the Enzcheck reagents to remove any contaminating phosphate that might convolute the experimental results.

